# Genome-wide landscape of RNA-binding protein dysregulation reveals a major impact on psychiatric disorder risk

**DOI:** 10.1101/2020.05.19.102319

**Authors:** Christopher Y. Park, Jian Zhou, Aaron K. Wong, Kathleen M. Chen, Chandra L. Theesfeld, Robert B. Darnell, Olga G. Troyanskaya

## Abstract

Despite the strong genetic basis of psychiatric disorders, the molecular origins of these diseases are still largely unmapped. RNA-binding proteins (RBPs) are responsible for most post-transcriptional regulation, from splicing to translational to localization. RBPs thus act as key gatekeepers of cellular homeostasis, especially in the brain. Here, we leverage a deep learning approach to interrogate variant effects genome-wide, and discover that the dysregulation of RBP target sites is a principal contributor to psychiatric disorder risk. We show that specific modes of RBP regulation are genetically linked to the heritability of psychiatric disorders, and demonstrate that diverse RBP regulatory functions are reflected in distinct genome-wide negative selection signatures. Notably, RBP dysregulation has a stronger impact on psychiatric disorders than common coding region variants and explains heritability not currently captured by large-scale molecular QTL studies (expression QTLs and splicing QTLs). We share genome-wide profiles of RBP target site dysregulation, which we used to identify DDHD2 as a candidate schizophrenia risk gene, in a public web server. This resource provides a novel analytical framework to connect the full range of RNA regulation to complex disease.

## Main

Interrogating the genetics behind psychiatric disorders is a key path in understanding the pathophysiological cause of mental illness, in part due to the largely immutable nature of the genome during the pathogenic manifestation. In particular, genome-wide associated studies (GWAS) have been the most widely adopted human genetics approach for studying psychiatric disorders. With ever-increasing sample cohorts, surpassing one million subjects for some psychiatric traits^1^, numerous risk loci have now been cataloged. Despite this progress, the pathological mechanisms and biochemical perturbations that lead to human psychiatric disorders are still lacking – a critical step for translating the genetic discoveries into actionable targets.

RNA-binding proteins (RBP) regulate various aspects of RNA metabolism, including RNA splicing^2^, localization^3^, stability^4^ and translation^5^. Each of these functions are critical not only in the expression of proteins, but in establishing their proper spatiotemporal function, especially in the brain^6^. Proper regulation of the full RNA life cycle by RBPs is particularly important in neurobiology, where complex regulatory events take place in synapses far away from the nucleus^7^.

RBP-encoding genes are frequently mutated in neuropsychiatric disorders, suggesting a pathogenic role^8–10^ and inspiring follow-up efforts to study the larger set of variants at the RNA targets that affect RBP-RNA interactions^11–15^. These studies have shed light on the roles of specific RBPs and functions, particularly splicing, in neurological disease. However, comprehensive and genome-wide insight into diverse RBPs, functions and causal target sites are lacking. Thus, to what extent dysregulated RNA-RBP interactions contribute to psychiatric disorders remains an open question.

Here, we address this challenge with the first genome-wide, systematic analysis of the role of RBP target site dysregulation in psychiatric disease. To map variant impacts on RBP-RNA interactions at scale, we leverage a deep learning-based sequence model, Seqweaver, whose accuracy in predicting variant effects on RBP target site dysregulation we previously extensively evaluated both computationally and experimentally, including by detecting *de novo* noncoding mutation signal in autism in probands versus their unaffected siblings^16^. We apply Seqweaver to profile allele-specific effects of inherited variants genome-wide, enabling us, for the first time, to examine the diverse landscape and impact of RBP dysregulation in complex psychiatric disorders.

Our study generates genome-wide variant level annotations of RBP dysregulation, which we make publicly available at hb.flatironinstitute.org/seqweaver. We show that the dysregulation of RBP target sites and diverse RBP regulatory functions are top drivers of psychiatric disorder risk. This analytical framework will accelerate biochemical investigation into disease genetics. Indeed, leveraging this resource as a case study, we discover a novel link between an RBP-disrupting variant in DDHD2, which encodes a phospholipase involved in hereditary spastic paraplegia, and a multiethnic-associated locus that increases the risk of schizophrenia.

## Results

### Regulatory function of RBPs is reflected in their genome-wide negative selection signature

Mutations that cause neuropsychiatric disorders are expected to decrease fitness, with epidemiology studies showing that these patients have reduced fecundity compared to the general population^17^. Consequently, we hypothesized that if RBP dysregulation contributes to brain-associated diseases, then variants disrupting RBP-RNA interactions should show a strong signature of negative selection.

To test this expectation, we leveraged the largest pool of human variants from control cohorts released by the Genome Aggregation Database (gnomAD)^18^. For each transcribed, noncoding variant in gnomAD (> 20 million SNPs), we interrogated the levels of RBP disruption, specifically using deep learning inference on 232 Seqweaver RBP models (RBP model list Supplementary Table 1). We found significant depletion of strong effect RBP-RNA disrupting variants at high allele frequencies (MAFs, p < 2.2 × 10^−16^ Wilcoxon rank sum test common vs ultra-rare). The mean variant RBP dysregulation effect increased significantly from common (MAF > 0.05) to ultra-rare inherited variants (MAF < 0.001, p < 2.2 × 10^−16^ Wald test, Supplementary Fig. 1), which is consistent with a major fitness impact and negative selection acting on variants that disrupt RBP target sites.

Within genes, RBPs frequently exert their regulation via noncoding regions, such as the 5’UTR, introns or 3’ UTR. We reasoned that the function of the RBP should be reflected in specific negative selection signatures in the relevant regions. For instance, RBPs that impact mRNA stability would show elevated negative selection signatures in 3’UTRs, whereas splicing RBPs would show these signatures in introns. Importantly, a differential selection signature would shed light on the function of RBPs, and thus providing complementary information to the biochemical studies of RBPs.

To define sub-genic selection signatures, we looked for statistically significant interactions between the genomic location of a variant and the degree of selection acting on RBP dysregulation. We found that diverse RBPs can be segregated by their selection signature across the 5’UTR, 3’UTR or introns (Fig. 1A, 212 RBP models with a Benjamini-Hochberg corrected FDR < 0.05 sub-genic annotation interaction, full results in Supplementary table 2). The significant association of each RBP with a sub-genic location rules out the baseline interpretation of confounding stochastic non-functional interactions, and points to an active regulatory function with specific fitness consequences.

**Figure 1.**
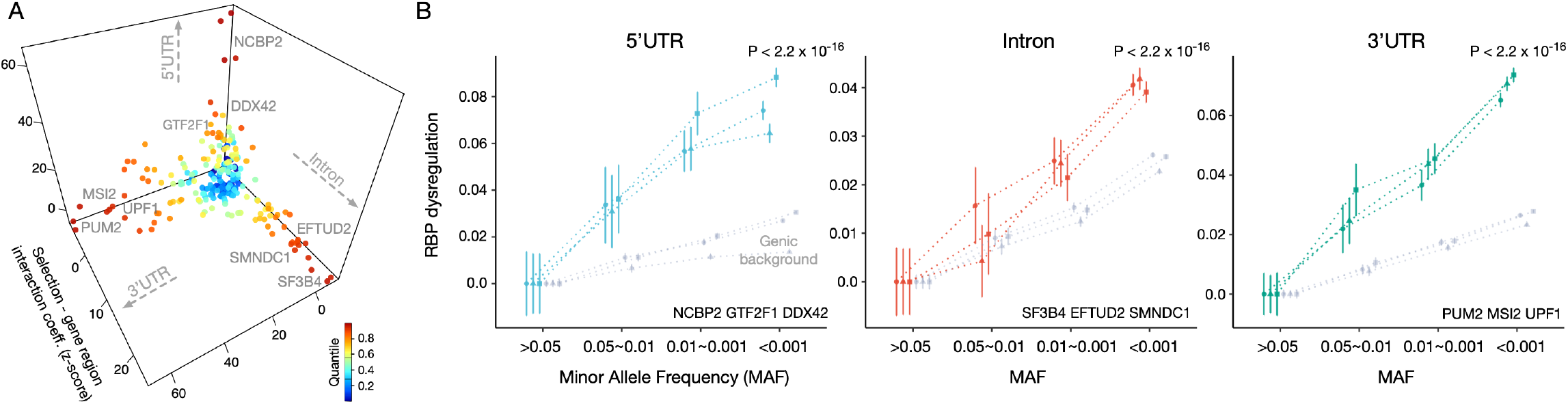
Negative selection signatures differentiate RBPs by their regulatory function. a) For each RBP, the negative selection signatures along the major axes of sub-genic regulatory regions (5’/3’ UTR and introns 200 bp flanking alternatively spliced exon) are plotted. Higher coefficient z-scores, along the x, y or z axes, implies stronger regional selection compared to the gene background. The fitness effects contributed by RBP regulatory roles, beyond splicing, are shown by the separation of RBPs along the regional selection axes. b) Examples of regional sub-genic selection signatures for the three major noncoding regions regulated by RBPs. GnomAD cohort noncoding variants (MAF bins x-axis) and variant set mean RBP dysregulation estimates (Y-axis) are shown. Stronger RBP selection signatures can be observed by the larger slope for each sub-genic region compared to the background gene levels. All inferred mean RBP dysregulation scores were normalized by subtracting average dysregulation predicted scores of common variants (>0.05 MAF) for comparison (95% CI).

For many known splicing factors (e.g. SMNDC1 and PRPF8; p < 2.2×10^−16^ Wald test on coefficient of interaction term), intronic variants displayed a significantly elevated level of selection spanning the spectrum of minor allele frequencies compared to the genic background (Fig. 1B). The cap-binding protein NCBP2 showed the most significant enrichment for 5’ UTRs, whereas for 3’UTRs, we confirmed the strong human fitness effect of many known mRNA stability, localization and polyA regulatory proteins (e.g. MSI2, PUM2, PABP and ELAVL; all p < 2.2×10^−16^).

Unexpectedly, we found that variants within the 3’UTR that disrupt binding of the superfamily I RNA helicase UPF1 are under a significantly elevated level of negative selection compared to the rest of the gene (p < 2.2 × 10^−16^). UPF1 is an essential component of the nonsense-mediated decay (NMD) machinery. Our finding is thus unanticipated, but consistent with recent biochemical reports that suggested that UPF1 binding to the 3’UTR can regulate target mRNA stability^19^. The regional selection enrichment of UPF1 provides strong corroborating genetic evidence for its role in 3’UTR-mediated post-transcriptional regulation, beyond its canonical NMD function.

### Variants that disrupt RBP binding is a major contributor to psychiatric disorder risk

Having established the strong importance of RBPs in selection and human fitness, we next investigated the potential role of RBP dysregulation on psychiatric disorder heritability. The strong family history of psychiatric disorders and high heritability estimates make this an especially important question^20,21^. To address this hypothesis, we applied the statistical framework of stratified LD score regression^22^ to partition disease heritability into various functional annotations while directly modeling the extensive LD structure between SNPs. The LD score regression framework allows estimation of SNP effects (*τ*\*) standardized for comparison across different disease or trait GWAS studies while conditioning on a collection of baseline functional annotations (e.g. coding region, allele age, CpG content, enhancers, promoter and epigenetic histone marks Methods)^23^. Here we combine LD score regression with our deep learning framework Seqweaver to estimate the contribution of RBP dysregulation to psychiatric disease.

The stratified LD score regression framework has been tested and shown to produce robust results in large collections of studies^22–24^. Nevertheless, we performed a comprehensive negative control test in the context of RBP dysregulation. We simulated genetic architecture traits where the underlying casual SNPs were sampled entirely from experimentally profiled brain enhancers, promotors, and brain-expressed protein coding regions (i.e. mostly non-RBP regulatory regions) using real genotypes from the 1000 Genomes Project^25^ (Methods). Overall, across the 232 RBP models, the simulations produced well-calibrated estimates of RBP dysregulation effect sizes without any upward bias (Supplementary Fig. 2), demonstrating the robustness of our regression models.

Having established the statistical framework, we focused on GWAS from five well-established polygenic psychiatric disorders: ADHD^26^, autism spectrum disorder^27^, bipolar disorder^28^, major depression^29^ and schizophrenia^30^. These GWAS were conducted with standardized analysis pipelines by the Psychiatric Genomics Consortium and therefore minimize the potential sources of technical artifacts. We observed significantly elevated levels of RBP dysregulation effect size (*τ*\*) estimates across all five psychiatric disorders, with 304 cases where dysregulation of targets of a specific RBP had a significant effect on a psychiatric disorder after correcting for multiple hypothesis testing (Fig. 2A, FDR < 0.05, Benjamini-Hochberg correction, Supplementary Table 3). These results demonstrate a strong underlying causal effect of RBP dysregulation in all five disorders (Fig. 2A).

**Figure 2.**
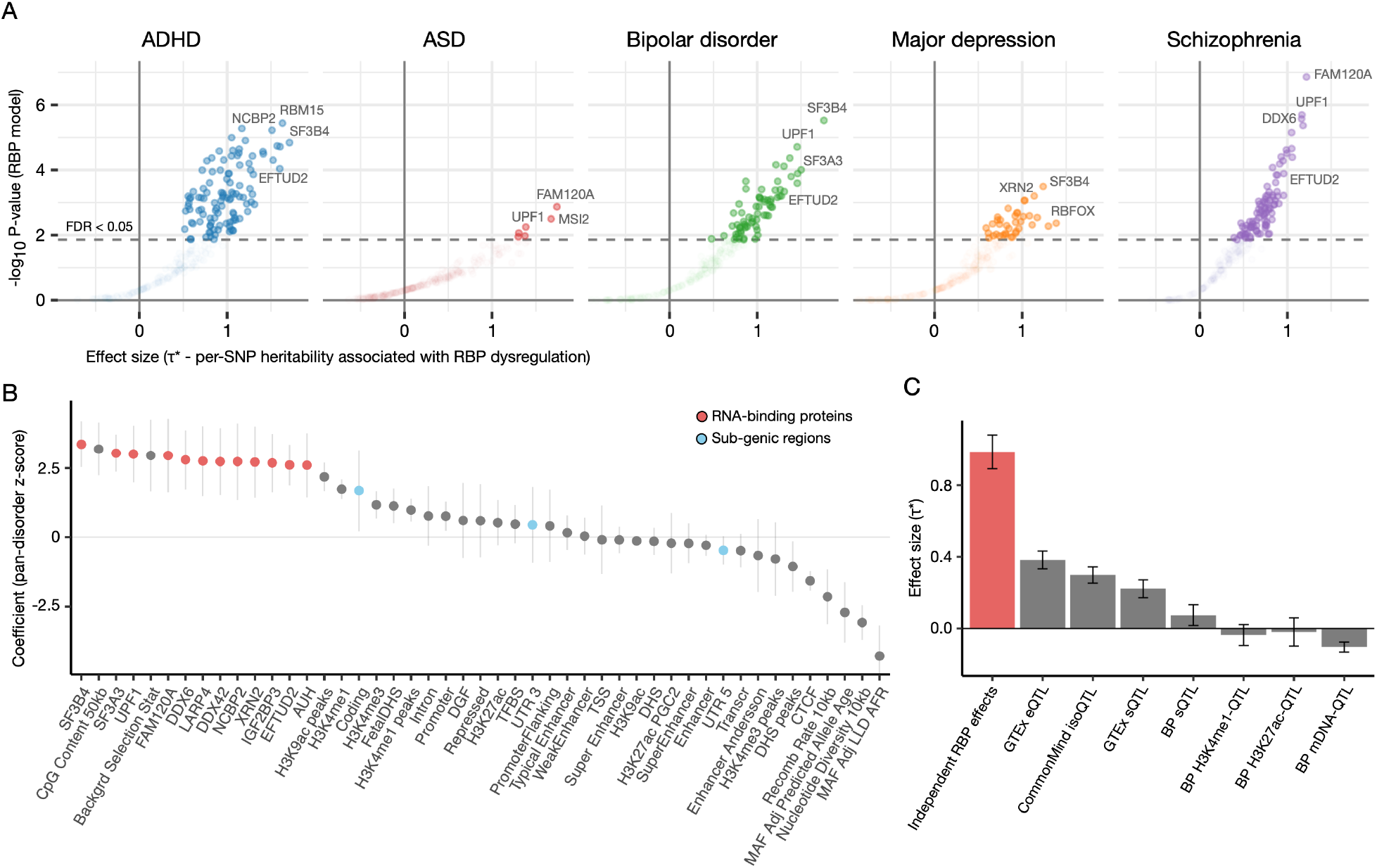
Genome-wide RBP dysregulation is a significant source of psychiatric disorder heritability. a) The per-SNP heritability effect sizes (*τ*\*) for each RBP target site dysregulation is plotted across the five major psychiatric disorders. The dashed line indicates RBP models below FDR 0.05 threshold after multiple hypothesis correction (block jackknife-based one-sided p-values; Benjamini-Hochberg correction). b) The statistical association between per-SNP heritability and the top psychiatric disorder-associated (TPA) RBPs (mean z scores across the five disorders). The jointly fit representative baseline annotations are also shown for comparison (sub-genic region annotations are highlighted blue e.g. coding region). c) The per-SNP heritability effect sizes (*τ*\*) for TPA RBPs after conditioning on a collection of molecular QTL annotations (i.e. independent RBP effects from molecular QTLs and baseline annotations). Genotype-Tissue Expression (GTEx), CommonMind, BLUEPRINT (BP). The jointly fit collection of QTL annotation effect sizes is also plotted. Expression (eQTL), splicing (sQTL), mRNA isoform (isoQTL), DNA methylation (mDNA QTL). All error bars are 95% CI.

In addition, we found novel associations between disrupted RBP target sites and RBPs that were previously associated with disease. For example, target sites of the spliceosome-associated factor EFTUD2 contributed significantly across psychiatric disorders (e.g. ADHD p=1.4×10^−4^, SCZ p=6.4×10^−4^ jackknife), and haploinsufficiency of EFTUD2 causes craniofacial malformation, microcephaly and developmental delay^9^, which are shared with many non-Mendelian neurological diseases. In addition, major depression was significantly associated with variants that disrupt target binding by RBFOX, a key splicing regulator in the brain^6^ (*τ*\*=1.4, p=8.6×10^−3^). Indeed, two GWAS loci within the RBFOX1 RBP gene locus itself are associated with major depression risk^29^. Overall these data suggest that psychiatric disease is significantly linked with perturbations not only of RBPs, but also their targets, which represent a much larger set of variants spread across the transcribed regions of the genome.

### RBP target effects exceed those of coding variants and broadly impact post-transcriptional regulation

Next, we examined how variants that dysregulate RBP function compare to other functional variant categories, by comparing across the jointly fit annotations in the regression models. We found that the statistical association between disease heritability and RBPs are among the top functional annotations, exceeding the collective set of coding variants or previously annotated epigenetic regions (Fig. 2B). Thus, noncoding variants that perturb RBP binding significantly increase psychiatric disorder risk. Furthermore, within gene regions, the collective liability of RBP dysregulation can exceed coding region variant effects (Supplementary Fig 3). These results remain significant and robust after conditioning on potential confounding factors, such as background selection rate, low levels of LD (LLD), allele age and minor allele frequency (Fig. 2B baseline annotations included in the regression model, Methods).

Notably, the strong noncoding dysregulation effects that we identified include diverse categories of RBPs, beyond splicing factors. For example, the most well-powered of the five GWAS, schizophrenia, 49/91 significant RBP models (FDR < 0.05) were UTR regulatory RBPs. This observation reveals a wider importance of post-transcriptional effects beyond splicing, and was not limited to schizophrenia: the top psychiatric disorder-associated (TPA) RBPs (mean z-score > 2.5) covered all the major RNA regulatory modes from splicing to transcript stability, based on the shared risk across all five psychiatric disorders studied (Fig. 2B). For example, UPF1^19^ and FAM120a^31^, which regulate transcript degradation, showed consistent strong signals across the disorders, with top ranked effect sizes in schizophrenia (UPF1 *τ*\*=1.16 p= 2.0×10^−6^, FAM120a *τ*\*=1.2 p= 1.4×10^−7^). We also observed a pair of ATP-dependent RNA helicases (DDX6, DDX42) among the TPA RBPs, adding support to the neuropathogenic role of helicases^32^. Overall, these data demonstrate that disruption of diverse types of post-transcriptional regulation are highly associated with psychiatric disease burden.

Finally, to replicate our findings of RBP dysregulation effects in an independent cohort, we utilized cross-psychiatric disorder GWAS summary statistics released by the iPSYCH consortium^33^. Risk variants identified in this study largely represent loci with shared underlying genetic risk factors across psychiatric disorders, and are based on a homogenous Danish population diagnosed using the same public healthcare system criteria. We focused this replication analysis on schizophrenia, which had no overlapping individuals within the Psychiatric Genomics Consortium and iPSYCH cohorts. We found highly significant concordance of RBP-associated risk between the two cohorts (P < 2.2 × 10^−16^ Wilcoxon rank sum test, see Supplementary Fig. 4 for RBP dysregulation effect sizes). Thus, our RBP dysregulation estimates are robust and not a by-product of population stratification.

### RBP target effects explain substantial heritability beyond known molecular QTLs

Previous reports have found that molecular QTLs are strongly enriched for disease heritability^13,34^. Thus we investigated whether the profiled RBP dysregulation effects capture information about disease that is independent of the large-scale molecular QTL studies. We estimated the effect size of each RBP for each disorder while jointly conditioning on the molecular QTL-based annotations from GTEx^24^, CommonMind^35^ and BLUEPRINT^36^ consortium (in addition to all baseline annotations). We found that TPA RBPs remain highly significant and display overall greater effect sizes compared to the QTL annotations (Fig. 2C, Supplementary Table 4). Importantly, this implies that RBP dysregulation effects are largely independent from known molecular QTLs and should provide an important additional tool for dissecting genetic architectures of disease.

### RBP dysregulation contributes to shared and distinct aspects of psychiatric disorders

Having established the significant role of RBP dysregulation in disease, we next examined if altered RBP-RNA interactions underlie the extensively shared genetic architectures across psychiatric phenotypes. At the gene level, psychiatric disorder heritability is enriched for mutation-intolerant genes^37^. Here we investigated if this enrichment could be in part explained by variants that impact RBP regulation. Indeed, we observed that RBP effect sizes were significantly larger for target site variants within loss-of-function (LoF)-intolerant genes (Fig. 3A, P < 2.2 × 10^−16^ paired Wilcoxon rank sum test for RBP effect size *τ*\*, LoF intolerant defined by ExAC^38^ and controlled for different baseline gene heritability enrichment levels Methods).

**Figure 3.**
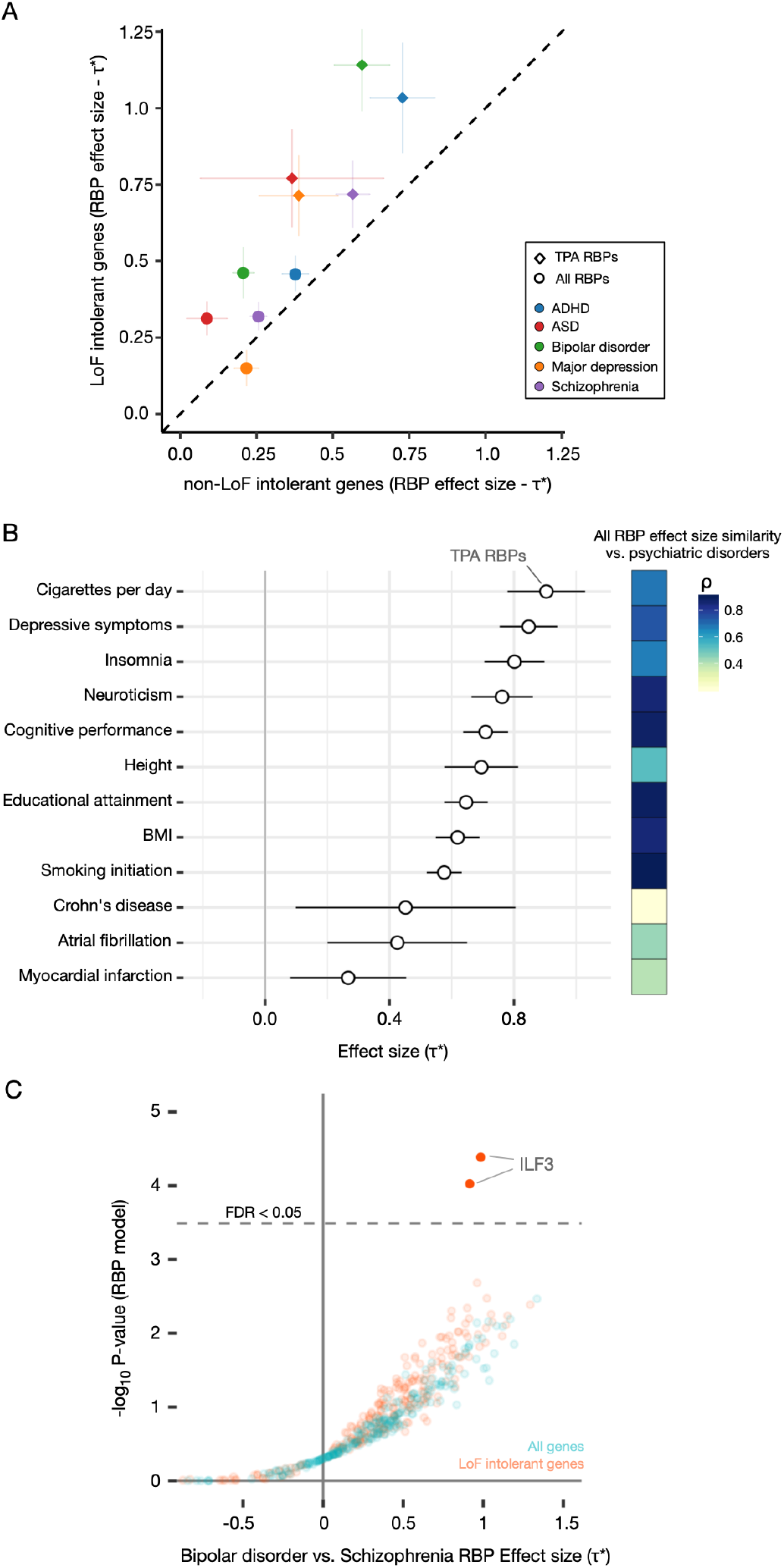
RBP dysregulation underlies shared and distinct genetic architectures of psychiatric phenotypes. a) The meta-analysis of RBP dysregulation effect sizes in loss-of-function (LoF) intolerant genes compared to the remaining set of genes is shown (95% CI). All effect sizes are conditioned on baseline annotations plus two additional annotations delineating LoF intolerant gene and their coding regions to control for the higher background heritability rates. b) Top psychiatric disorder-associated (TPA) RBP estimated effect sizes across a collection of psychiatric traits and non-brain related phenotypes are shown (95% CI). The adjacent heatmap displays spearman rank correlation for all RBP effect sizes between the phenotype and psychiatric disorders (mean effect). c) RBP dysregulation effect sizes (*τ*\*) for differential risk between bipolar disorder and schizophrenia, estimated for both all genes and for LoF intolerant genes. The dashed line indicates RBP models below FDR 0.05 threshold (Benjamini-Hochberg method).

Moreover, we found that numerous RBP models were significantly associated with heritability for a broader set of psychiatric and related phenotypes examined by the Brainstorm Consortium^24^ (Supplementary Fig. 5, 856 RBP model-phenotype pairs FDR < 0.05 with Benjamini-Hochberg correction). This association was especially strong for psychiatric traits like “Cigarettes per day” (a common proxy for addictive risk-taking behavior^39^) and “Depressive symptoms” (a widely shared clinical feature for many psychiatric disorders^40^), whereas the non-brain related phenotypes displayed lower correlation of overall RBP effect sizes with psychiatric disorders (Fig. 3B). These results imply that variants disrupting RNA-RBP interactions affect neuropathogenic pathways and are a significant driver of the high genetic correlations observed between psychiatric disorders and traits.

On the other hand, we found that the disruption of distinct RBP target sites can help explain differences between psychiatric disorders. Specifically, we found that disruption of targets for the RBP ILF3 contributes to the differential liability between schizophrenia and bipolar disorder (Fig. 3C, two biological replicate ILF3 models highlighted, p=4.1×10^−5^ LoF intolerant genes, jackknife), extending findings of the Psychiatric Genomics Consortium study^28^. Moreover, at the gene-level, ILF3 was the 5^th^ most significantly associated gene in bipolar disorder (Supplementary Fig. 6A, p=1.2×10^−9^, MAGMA statistical framework^41^, colocalization analysis further supports this ILF3 association Supplementary Fig. 6B), but it had no significant association in the better-powered schizophrenia study^30^. Our analysis suggests that the molecular network composed of both RBP ILF3 and its targets helps to discern these two psychiatric disorders. Determining how the ILF3 network alters brain cellular functions will shed light on how genetics influence clinical outcome variation.

### Functional mapping based on RBP dysregulation identifies DDHD2 as a schizophrenia risk gene

Hundreds of genomic regions are associated with the risk of psychiatric disorders, consistent with a polygenic architecture^30^. However, very few disease-associated regions have been mapped to their causal SNPs, with the underlying biochemical mechanism dissected. As a case study, we leveraged our ability to interrogate allelic-specific RBP target site dysregulation genome-wide to investigate a schizophrenia risk region.

The 8p12 genomic region was first identified as a significant schizophrenia risk region in the Han Chinese population^42^ and was subsequently found significantly linked in Europeans^37^. Cross-population replication implies a robust molecular cause underlying the associated loci with global clinical prospective. Our analysis provides a potential biochemical mechanism for this association: within this region we identified a SNP in the DDHD2 3’UTR that can disrupt binding by the RBP QKI, which is known to play an important role in schizophrenia^43,44^ (Fig. 4A, this top Seqweaver predicted SNP rs6981405 was a fine-mapped candidate SNP (95% credible set)^37^).

**Figure 4.**
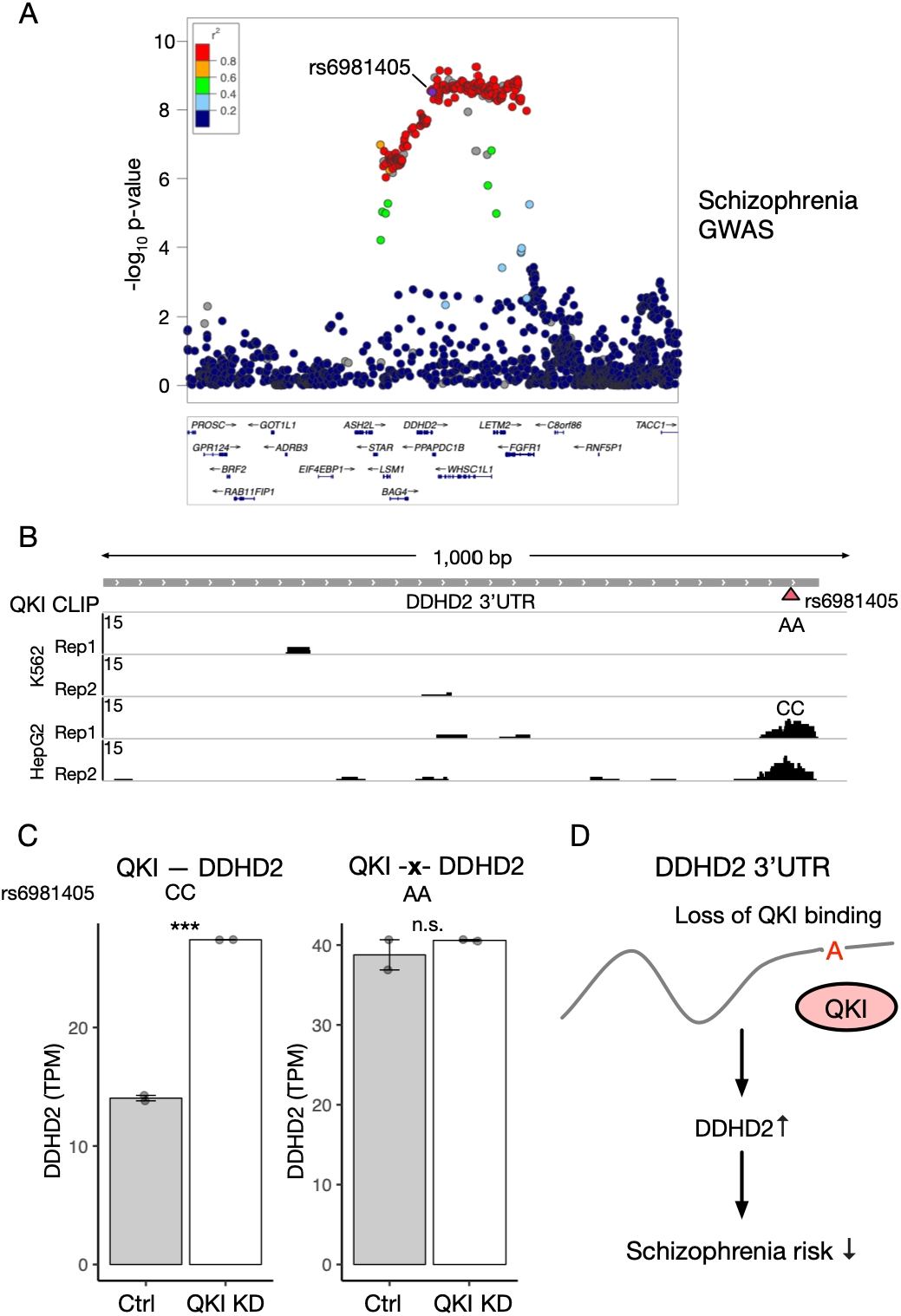
Functional RBP regulatory mapping identifies a schizophrenia risk variant in DDHD2 3’UTR. a) Schizophrenia GWAS signal for the cross-ethnic associated region. The highlighted SNP rs6981405 represents the top predicted RBP dysregulation variant disrupting RBP QKI binding. ENCODE eCLIP data confirms SNP rs6981405 C>A leads to the disruption of RBP QKI binding to DDHD2 (rs6981405 genotype for cell line K562 homozygous AA, HepG2 homozygous CC). SNPs were allowed during CLIP read alignment. c) QKI knockdown coupled by RNA-seq confirms the QKI-mediated regulation of DDHD2, which is disrupted in the homozygous AA genotype (i.e. QKI KD shows no effect when SNP rs6981405 impedes with RBP binding). Error bars represent SEM. d) Schematic: the variant at rs6981405 disrupts the QKI - DDHD2 3’UTR interaction, which alters the abundance of mature DDHD2 mRNA, and, in turn, schizophrenia risk.

ENCODE^45^ QKI eCLIP data, available in two cell lines (K562 and HepG2), support this association. The candidate SNP rs6981405 is homozygous CC in HepG2 and homozygous AA in K562. QKI and its target DDHD2 are highly expressed in both cell lines (> 15 TPM). Importantly, QKI-DDHD2 binding is observed only in homozygous C allele genotype (Fig. 4B), consistent with our estimation (i.e. A allele disruption of QKI binding). Furthermore, RNAi-mediated depletion of QKI led to elevated levels of DDHD2 mRNA in C allele genotype, but not in homozygous A allele genotype line, where QKI binding is already disrupted (Fig. 4C). Thus, mutation of this SNP in DDHD2 mRNA precludes its regulation by QKI.

DDHD2 is a principal brain triglyceride lipase, that when mutated causes a hereditary neurological disease, spastic paraplegia^46^. Our genetic evidence, provided by RBP regulatory mapping, coupled with supporting experimental data, suggest that QKI-mediated regulation of DDHD2 transcript levels influences the risk of schizophrenia and imply a pathogenic role for altered lipid metabolism in this disease.

## Discussion

A critical challenge in human disease research involves moving from cataloging disease risk loci to understanding the molecular mechanism. Interactions between transcripts and RBPs are early events that influence protein expression and function. Therefore, interrogating genetic architectures at this early layer of molecular regulation presents a powerful opportunity, as it reduces the complexity of having to deconvolute disease correlated signal into causal factors that often burdens the analysis for further downstream approaches. Most importantly, targeted biochemical perturbation of RBP-RNA interactions has proven to be an effective new avenue of clinical intervention^47^, and therefore delineating RBP target sites as a major source of molecular dysfunction contributing to psychiatric disorders is a vital task.

In this work, we establish that RBP dysregulation is a key contributor to human fitness by identifying extensive negative selection signatures in the largest-to-date WGS gnomAD^18^ cohort. We further find that concentrated regional fitness effects observed for each RBP provide a genetic indicator for the underlying biochemical regulatory function. We also highlight that disruption of diverse RBP functions significantly affect fitness, supporting an extensive pathogenic contribution beyond splicing regulation.

Next, focusing on psychiatric disorders, we provide support for a significant causal role for RBP dysregulation, linking inherited risk variants to biochemical perturbations that ultimately lead to psychiatric clinical phenotypes. Intriguingly, one key theme we find that emerged is the psychiatric risk convergence at both the RBP regulatory factor (i.e. RBP protein itself) and its downstream target site dysregulation. For instance, variants within RBFOX and its downstream targets are linked to major depression risk, and variants within EFTUD2 and its downstream targets are linked to neurological dysfunction. In addition, we find converging evidence links RBP ILF3 and its RNA targets to the molecular differences between schizophrenia and bipolar disorder. Similarly, the RBP “fragile X mental retardation protein” (FMRP) is the most common monogenic cause of autism^48^, and FMRP mRNA targets are highly linked to both autism and schizophrenia^49–51^. Thus, these converging RBP regulatory networks may present ideal clinical targets, due to the greater collective biochemical contribution to pathogenicity.

Methodologically, we demonstrate that our deep learning inference of genome-wide molecular effects allows us to estimate major modes of biochemical perturbation and their contribution to disease. We find that splicing disruption is the tip of the iceberg, as we discovered widespread psychiatric disease risk associated to RBPs that regulate each step of the RNA life cycle from processing to localization to degradation. Current molecular QTL resources, while incredibly valuable, lack the breadth to capture these diverse molecular functions of RBPs. This caveat limits the scope of analysis for disease, now encapsulating hundreds of thousands of cohort samples (e.g UK biobank^52^). Indeed, there has been increasing evidence of an extensive pathogenic role of RBPs from cancer^53^, autoimmune disease^54,55^ to myopathy^56^. Our computational framework can enable studies of RBP dysregulation in these and other disorders on the whole-genome-scale. Furthermore, as more tissue- and cell-type-specific CLIP data becomes available, this approach can provide a data-driven window into tissue-specific RBP dysregulation in disease.

To enable rapid analysis of psychiatric diseases and the extension to the greater collection of disease GWAS studies, we now profiled and made available genome-wide inference of RBP target site dysregulation effects for the largest collection of human variation identified by the gnomAD cohort. This resource, with the entire spectrum of common to ultra-rare variants, should provide the means to interrogate RBP-derived human diseases at an unprecedented scale.

## Methods

### Deep learning inference of RBP dysregulation variant effects

We utilized the machine learning approach of deep convolutional neural networks (CNN)^57^ to build a quantitative model of the RNA sequence features required for each RBP binding biochemically assayed by CLIP^58^ (training data). These RBP models subsequently enable the probabilistic inference of variant impact, capturing both direct and indirect effects, on the RBP binding potential. The applied Seqweaver RBP model architecture and training are described in our previous *de novo* mutation autism work^16^, and to ensure that our current results can be directly comparable, we used the identically 232 CLIP-based RBP models (Supplementary Table 1) without any modifications (i.e. no additional training or manual parameter changes).

Briefly, CNNs allow researchers to design network architectures that can leverage information of high order motifs at different spatial scales but with optimal parameter sharing to avoid overfitting. Our Seqweaver DNN architecture consists of an initial input layer followed by a series of convolution and pooling layers. The input sequence layer contains a 4 × 1,000 matrix that encodes the input RNA sequence of U, A, G, C across the 1,000 bp window anchored around the RBP binding site. The subsequent convolution layer looks at 8 bps at a time shifting by 1 bp and computes the convolution operation of 160 kernels. At this first convolution level, the kernels are equivalent to searching for a collection of local sequence motifs in a one-dimensional RNA sequence. Analogues to neurons, we then apply a rectifier activation function (ReLU) that sets the convolution layer output to a scale of minimum of 0 (i.e. ReLU(x) = max(0,x)). Thus formally, input *S* results in convolution layer output location *n* for kernel *k* as the following:

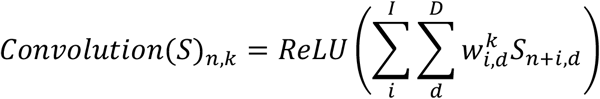

where *I* is the window size and *D* is the input depth (e.g. for the first convolution layer *D* represents the four RNA bases).

Next, we add a pooling layer that allows the reduction of the dimensional size of the network and parameters. Specifically, every window of 4 for a kernel output are collapsed into the maximum value observed in that span. Subsequently, the resulting output is used as input for a sequence of convolution (2^nd^), ReLU, pooling and convolution layer (3^rd^) in which higher order sequence motifs can be derived based on the first layer local motifs (2^nd^ conv. layer 320 kernels, 3^rd^ conv. layer 480 kernels with identical ReLU and pooling layer).

Lastly, we add a fully connected layer that can now take the resulting output from the three convolution steps to integrate across the entire 1,000 bp context to derive a final set of higher order sequence motifs. These sequence motifs are shared across all RBP models that allow optimal parameter reduction, but also are based on the biological intuition that many RNA sequence features are shared in the cell (e.g. RNA polyA signal, splice sites and branchpoints). The fully connected layer outputs are then subjected to RBP-specific weighted logistic functions (sigmoid, [0,1] scale) allowing for the simultaneous prediction of each RBP binding potential to the input RNA sequence.

Finally, for variant effect prediction, we take the absolute predicted probability differences between the two alleles (reference vs. alternative) computed by the convolutional neural network for each 232 RBP models. Important to note that no variant-level sequence information was used during the training of our Seqweaver RBP models, therefore we are not limited or biased by any variant-level training set. The final RBP variant effects were set to [0,1] scale for fitting LD score regression models. All training data and RBP models can be downloaded at hb.flatironinstitute.org/Seqweaver.

### Genome-wide RBP dysregulation analysis of negative selection

The 2.1 release of gnomAD^18^ cohort variants, all passing the random forest gnomAD quality filter were downloaded and filtered for noncoding region variants (i.e. nonrepeat regions of 5’UTR, intron and 3’UTR) of protein coding genes (we use AC >1 to filter for inherited variants). The resulting final 21,513,861 SNP variants were used in the analysis.

For each Seqweaver RBP model, the distribution of absolute predicted probability differences (ref. vs alt. allele) across all variants were standardized to have a standard deviation 1 to obtain the final RBP dysregulation estimates. The gnomAD cohort allele frequencies were used to segregate the variants into four different minor allele frequency bins (>0.05, 0.05~0.01, 0.01~0.001, <0.001) and then to obtain the mean variant RBP dysregulation levels.

Variant level annotations to sub-genic regions were conducted as previously described^16^, annotating to 5’UTR, 3’UTR or 200bps introns flanking an exon previously observed to be alternatively spliced^59^. The RBP sub-genic selection signature was assessed by fitting a linear model regressing RBP dysregulation levels on allele frequency and sub-genic annotations, and evaluating the statistical significance of the interaction term between the two explanatory variables. That is querying for statistically significant interactions between a variants gene location and the degree of selection acting on RBP target site dysregulation. For downstream analysis, UTR regulatory RBPs were defined by RBP models that showed sub-genic selection signatures with FDR < 0.05 for only 3’ or 5’ UTR regions.

### Estimating the RBP dysregulation GWAS effect sizes

The extensive linkage disequilibrium (LD) between SNPs in the human population provide an analytical challenge for estimating the true underlying effect size for RBP dysregulation from GWAS. For example, high *χ*^2^ statistic SNPs in the 3’UTR may appear to be an indication of UTR mediated risk to a disease, but conversely may simply just be tagging the true causal enrichment of protein coding region SNPs due to the high LD in the region. To resolve this challenge, we use the previously published statistical framework of stratified LD score regression^22^ to estimate the RBP dysregulation effect sizes for each examined trait or disease GWAS. More specifically from the summary statistics of a GWAS, we can write the expected *χ*^2^ value for SNP j as

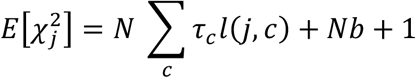

where *N* is sample size and the annotation specific “LD score” *l*(*j*, *c*), representing annotation (*c*)’s cumulative effects tagged by the SNP j, can be written as

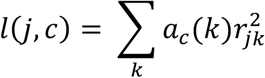

where *a_c_*(*k*) is the annotation value at SNP k (e.g RBP dysregulation level or coding SNP), *r*_*jk*_. is the correlation between SNP j and k in the reference panel (selected to best match the GWAS cohort), and b measuring the confounding bias^22^. Lastly, *τ*_*c*_ and the final standardized form 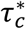 – normalized by the total SNP-based heritability and s.d. of an annotation – represents the estimated effect size of the annotation^23^.

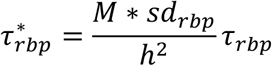

More formally for RBP dysregulation annotations, *τ*\* represents the per-SNP heritability (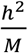, *M* number of common SNPs) associated with a standard deviation increase of variant RBP effect (*sd_rbp_*). We restrict our RBP predictions to SNPs from 1000 Genomes Project (European cohort), and fit the regression model only on HapMap SNPs with MAF > 0.05 as previously conducted^22^. Block jackknife procedure is used to test the statistically deviation from zero for each fitted *τ*\*.

As presented in the regression model, we fit 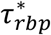 by conditioning on a collection of baseline annotations to avoid upward bias in the effect size estimation. We obtain the collection of baseline annotations previously used in the stratified LD score regression study (i.e. baselineLD)^23^, that includes functional annotations such as coding regions, 3’/5’ UTR, intron, promoter, transcription start site (TSS) and multiple epigenetic marks. We included a new functional baseline annotation that labels all gene region SNPs, controlling for baseline effects tagging transcribed regions, that collectively results in appropriately calibrated null uniform RBP p-values based on permutation test shown in Supplementary Fig. 7. Additional baseline annotations include non-functional annotations such as allele age, minor allele frequency, low levels of LD, CpG content and background selection statistics. We excluded conservation-based annotations, as RBP regulatory binding sites are known to be highly conserved^60–62^. The final reported RBP effect size 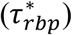 were jointly fit, iteratively for each RBP, with all baseline annotations (full 71 baseline annotations listed in Supplementary Table 5).

### Simulations for RBP effect size estimation

We conducted simulations to ensure that our regression models produced unbiased RBP effect sizes. Specifically, we tested to verify that false positive results were not obtained for genetic architectures where the causal SNPs were derived from functional elements that are largely non-RBP regulatory regions – epigenetic enhancers, promoters and protein coding regions. We simulated 400 GWAS summary statistics using the 1,000 Genomes Project European reference panel using simGWAS^63^. Testing for two scenarios, in each simulation, we sampled 1% or 5% SNPs (MAF > 0.01 and chromosome 1), as the causal set from brain epigenetic enhancers annotated by the PsychENCODE Consortium^64^, and both promotors^65^ and coding regions (restricted to nonsynonymous variants) that are expressed in the brain^66^. For each causal SNP effect size, we model as a Fisher polygenic model with trait heritability set to *h*^2^ = 0.5. Each simulated GWAS was fit with our LD score regression model (RBP + baseline annotations) to obtain the RBP effect size estimate 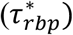. Results for the simulations produced overall robust unbiased estimates across our RBP models (Supplementary Fig. 2).

### GWAS disease and trait selection

We selected psychiatric disorder GWAS studies conducted by the Psychiatric Genomics Consortium, that was uniformly processed and analyzed, and was sufficiently powered to observe genome-wide significant SNPs, resulting in five disorders – ADHD, autism spectrum disorder, bipolar disorder, major depression and schizophrenia. To facilitate cross-study comparison, we selected psychiatric traits and non-brain associated diseases previously profiled by the Brainstorm consortium study (excluded traits that did not find genome-wide significant SNPs). The Danish cohort from iPSYCH consortium cross-disorder GWAS study^33^ was used as replication analysis. Population matched schizophrenia case vs. bipolar disorder case GWAS^28^ summary statistic was obtained from the Psychiatric Genomics Consortium web portal (full list of GWAS studies examined in this work Supplementary Table 6).

### Joint modeling of molecular QTLs

Fine-mapped GTEx^24^ eQTL (FE-meta) and BLUEPRINT^45^ molecular QTL annotations were obtained from a previous study^34^, that generated and validated the max causal posterior probability (MaxCPP)-based QTL annotations for GWAS enrichment analysis. The CommonMind^67^ isoformQTLs and GTEx sQTL (brain cortex version 8) were fine-mapped to produce MaxCPP annotations following the same procedure as was previously reported^34^. The collection of molecular QTL MaxCPP annotations and all baseline annotations were jointly modeled in the stratified LD score regression with each RBP annotation to estimate the disease associated effect sizes.

### Genetic architecture analysis

Loss-of-function intolerant genes were obtained by the ExAC consortium with pLI threshold of > 0.9 as previously described^16^. For stratified LD score regression models, we jointly fit, for each RBP model, the LoF intolerant gene and non-LoF intolerant gene regions variant RBP effect sizes (*τ*\*) by splitting the RBP annotation into two by gene set. We added two additional baseline annotations for this analysis, that includes SNP to LoF intolerant gene regions, and SNP to LoF intolerant gene’s coding region, to prevent potential upward bias due to the general higher background heritability enrichment levels. We also added the two SNP to LoF intolerant gene or their coding region baseline annotations for the differential risk analysis between schizophrenia and bipolar disorder to mitigate any potential bias.

MAGMA^41^ was used to estimate the gene level association with schizophrenia and bipolar disorder. GENCODE^68^ v25 gene annotations lifted to GRCh37 coordinates and total 19,984 protein coding genes were analyzed. We used SNPs from upstream 10k, gene body and downstream 1.5k for each gene as was previously used in a Psychiatric Genomics Consortium schizophrenia GWAS analysis study^69^ along with the 1000 Genomes Project^25^ European reference panel. Colocalization analysis for the ILF3 locus between schizophrenia vs. bipolar GWAS^28^ and GTEx ILF3 eQTL (v8 meta-tissue) was conducted using Coloc^70^.

### Functional mapping of DDHD2

QKI eCLIP and knockdown RNA-seq data was obtained by the ENCODE project^45^ in K562 and HepG2 cell lines. eCLIP data was processed as previously described^16^ and visualized in IGV^71^. Kallisto^72^ coupled with Sleuth^73^ was used for differential expression analysis of DDHD2 transcript (ENST00000520272) following QKI KD. P-values were computed using likelihood ratio test implemented in Sleuth and FDR was computed across all transcripts using Storey’s q-value method^74^. Genotyping results for SNP rs6981405 in K562 and HepG2 lines were obtained from ENCODE project.

## Code and data availability

The code is available from https://hb.flatironinstitute.org/seqweaver/about. All variant predicted scores have been made available to download and as an interactive web interface available at https://hb.flatironinstitute.org/seqweaver.

