## Supplemental Information for "Genome-wide landscape of RNA-binding protein dysregulation reveals a major impact on psychiatric disorder risk"

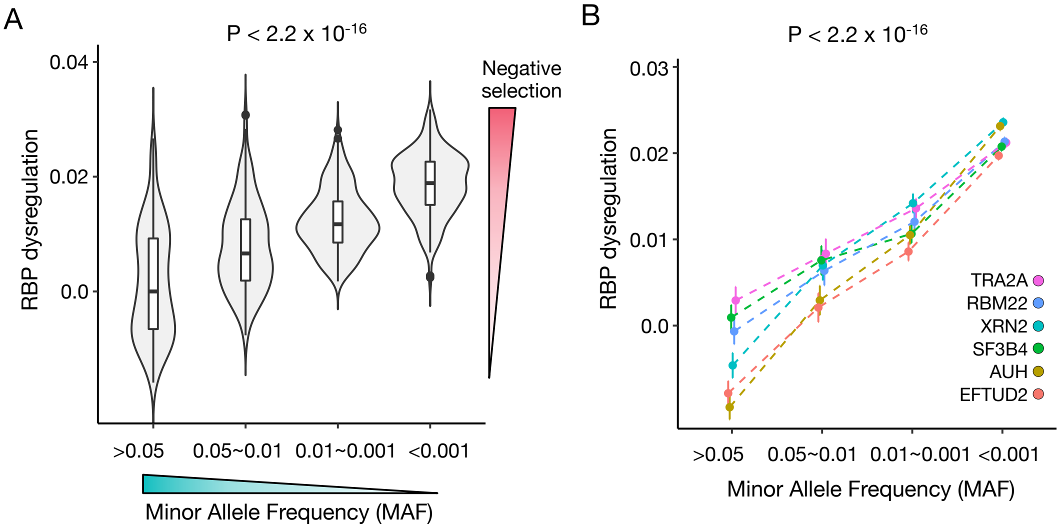


**Supplementary Fig 1. Population genetics revels negative selection acting on RBP target site dysregulation** a) Across the Seqweaver profiled RBPs, we observe differential selection signatures for variants when segregated by their RBP target site dysregulation levels. Specifically, for gnomAD cohort noncoding variants (MAF bins x-axis), mean RBP dysregulation (Y-axis) shows an inverse relation with allele frequency, consistent with significant negative selection acting on high impact RBP disrupting variants. b) The top RBPs previously implicated by their autism *de novo* mutation risk (Zhou, Park, Theesfeld et al.), all show significant negative selection signatures, consistent with selection impeding RBP impacting variants from reaching high population prevalence. P-values from Wald test for slope and all inferred mean RBP dysregulation scores were normalized by subtracting average dysregulation predicted scores of common variants (MAF > 0.05) for comparison (95% CI).

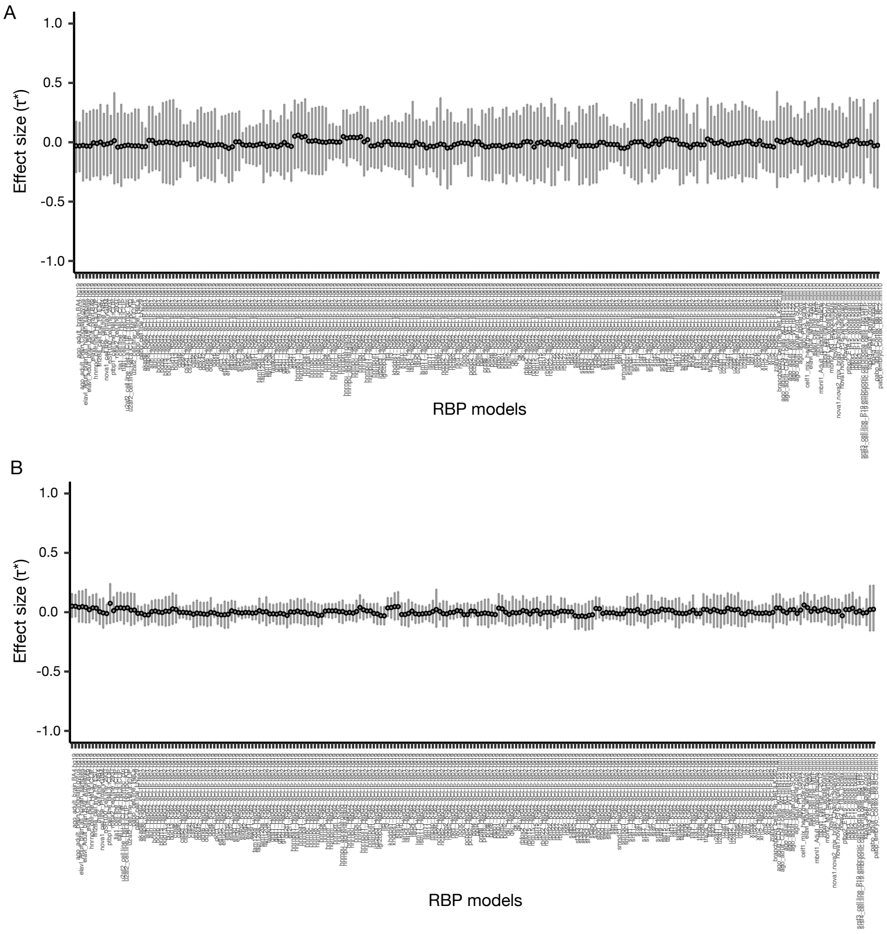


**Supplementary Figure 2. Simulation test confirm robustness of RBP effect size estimates.** To test that our regression models produce unbiased estimates of RBP effects, we simulated 400 GWASs with causal SNPs sampled from brain enhancers, promotors and nonsynonymous variants from brain expressed genes – representing mostly non-RBP regulatory regions (sample rate (A) 1% or (B) 5% ). The standardized RBP effect size estimates ($\tau^*_{rbp}$), jointly fit with baseline annotations, produced overall unbiased estimates across our RBP models. The median and the interquartile range of fitted RBP effect sizes are plotted.

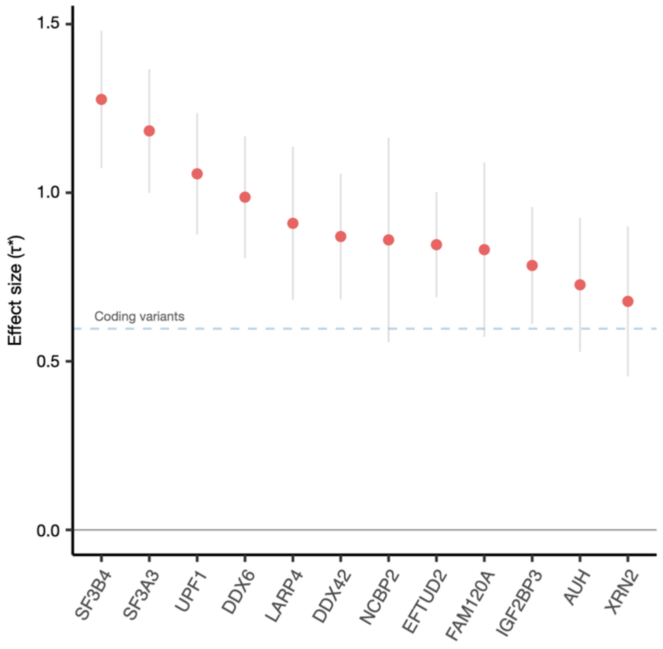


**Supplementary Figure 3. TPA RBP dysregulation effects compared to coding regions.** The mean effect sizes ($\tau^*$) across the five psychiatric disorders (ADHD, autism spectrum disorder, bipolar disorder, major depression and schizophrenia) for the top psychiatric disorder-associated (TPA) RBPs are plotted in comparison to common coding region pan-disorder variant effects (90% CI).

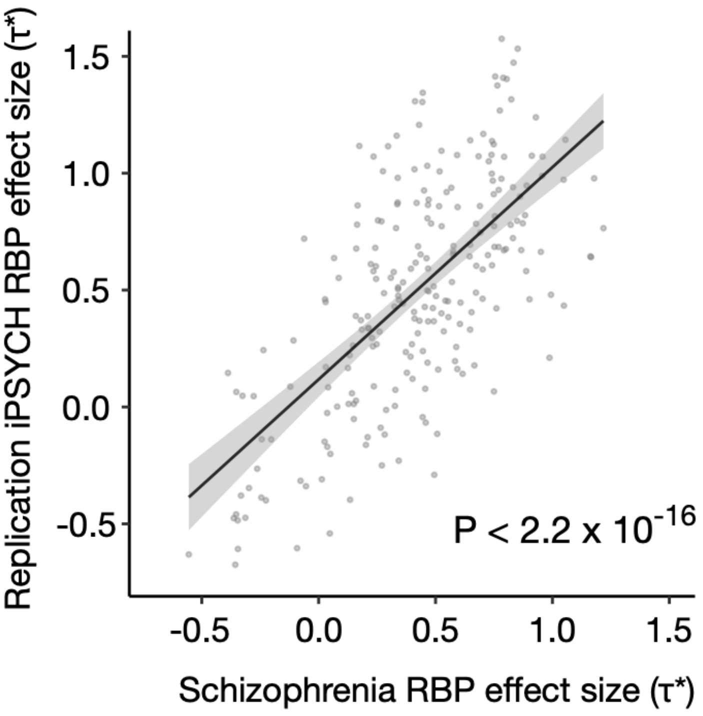


**Supplementary Figure 4. RBP dysregulation effects replicate in an independent cohort.** Replication of estimated schizophrenia RBP target site dysregulation effect sizes in the iPSYCH cohort ($\tau^*$). P-value computed using Wilcoxon rank sum test of RBP effect sizes.

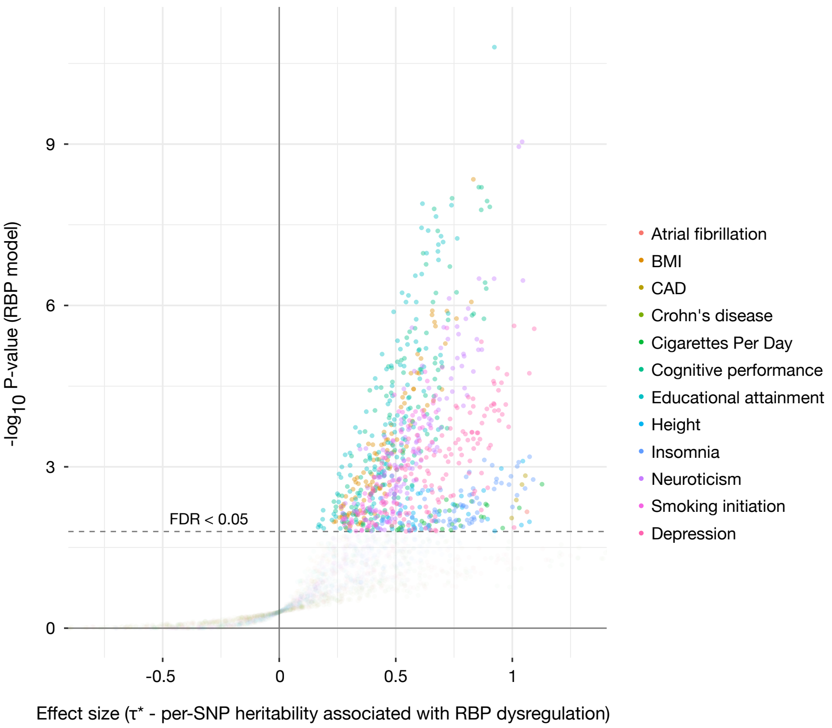


**Supplementary Figure 5. RBP dysregulation is a major contributor to human phenotypic variation.** The per-SNP heritability effect sizes ($\tau^*$) for each RBP dysregulation is plotted across a collection of psychiatric traits, brain-associated anthropomorphic traits and representative non-brain related phenotypes previously examined by the Brainstorm Consortium study. The dashed line indicates RBP models below FDR 0.05 threshold after multiple hypothesis correction (block jackknife-based one-sided p-values; Benjamini-Hochberg correction).

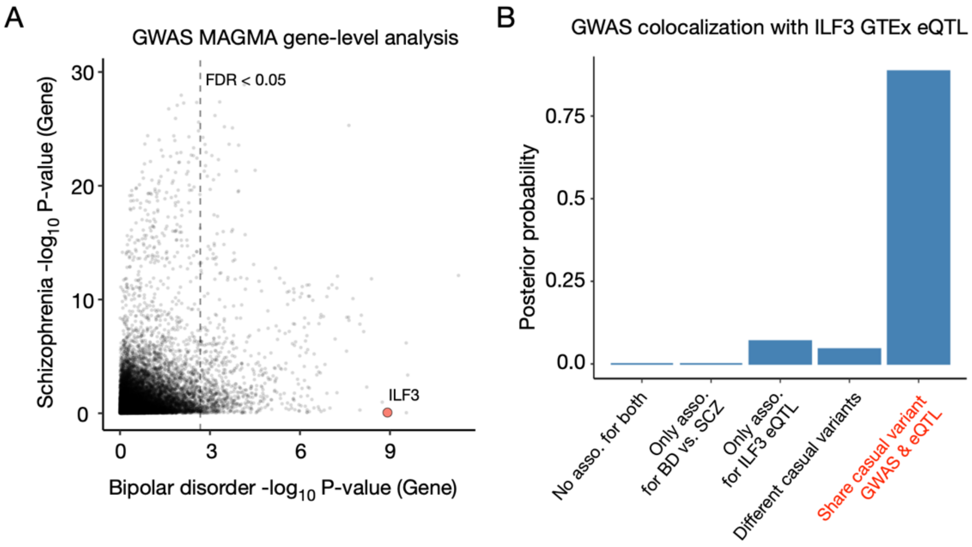


**Supplementary Figure 6. RBP ILF3 contributes to the differential liability of bipolar disorder vs. schizophrenia.** a) MAGMA gene locus GWAS association analysis shown for bipolar disorder and schizophrenia. Highlighted dot represents the RBP ILF3 gene locus statistical association in each of the two disorders. b) Colocalization analysis links the bipolar disorder case vs. schizophrenia case GWAS ILF3 gene locus association to ILF3 mRNA transcript levels (GTEx eQTL FE-Meta).

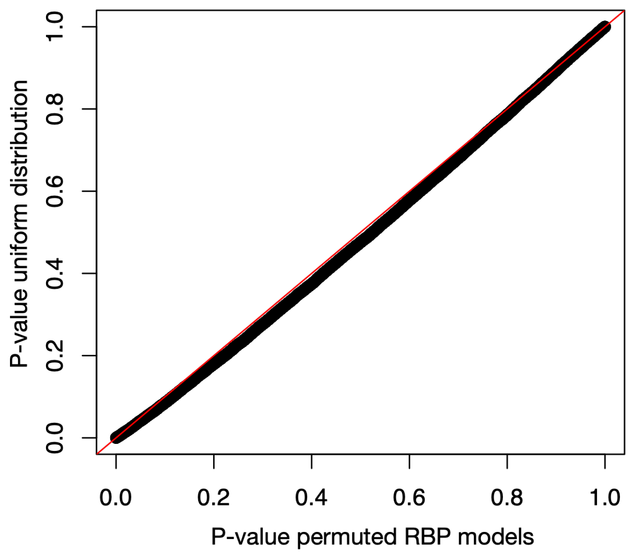


**Supplementary Figure 7. Permutation test of RBP effect size p-values.** We permuted the non-zero RBP target site dysregulation scores (i.e. gene region SNPs) and estimated the stratified LD score coefficient p-values for each RBP model across the five psychiatric disorder GWASs. With the inclusion of an additional baseline annotation mapping SNPs to gene, and therefore removing the implicit tagging of transcribed genomic regions, we find that our null p-value distribution is properly calibrated.
